# Proteasomal downregulation of the pro-apoptotic MST2 pathway contributes to BRAF inhibitor resistance in melanoma

**DOI:** 10.1101/2022.02.21.481296

**Authors:** David Romano, Lucia García-Gutiérrez, David J. Duffy, Keith T. Flaherty, Dennie T. Frederick, Walter Kolch, David Matallanas

## Abstract

The RAS-RAF-MEK-ERK pathway is hyperactivated in almost all malignant melanomas, and mutations in BRAF or NRAS account for most of these cases. BRAF inhibitors are highly efficient for treating patients with BRAF^V600E^ mutations, but tumours frequently acquire resistance within a few months. Multiple resistance mechanisms have been identified, due to mutations or network adaptations that revive ERK signalling. We have previously shown that RAF proteins inhibit the MST2 proapoptotic pathway in a kinase independent fashion. Here, we have investigated the role of the MST2 pathway in mediating resistance to BRAF inhibitors. We show that the BRAF^V600E^ mutant protein, but not the wildtype BRAF protein, strongly binds to MST2 and inhibits MST2 pro-apoptotic signalling. Downregulation of MST2 reduces BRAF inhibitor induced apoptosis. In BRAF inhibitor resistant cell lines MST2 pathway proteins are downregulated by ubiquitination and subsequent proteasomal degradation rendering cells refractory to MST2 pathway induced apoptosis. Restoration of apoptosis can be achieved by increasing MST2 pathway protein expression using proteasome inhibitors. In summary, we show that the MST2 pathway plays a role in the acquisition of BRAF inhibitor resistance in melanoma.

## Introduction

Metastatic melanoma is the most aggressive form of skin cancer, and its incidence is increasing worldwide, in particular in western countries (Rigel, 2010, Feigelson et al., 2019, Davey et al., 2021). This tumour type is characterised by a high frequency of genetic mutations within the RAS-RAF-MEK-ERK pathway, with mutations in BRAF, NRAS and NF1 as the main drivers of transformation (Curtin et al., 2005). Development of targeted therapies to rescue the effect that aberrations in these genes have in melanocytes has been the focus of intense research. This led to the development of BRAF inhibitors (BRAFi), which are extremely effective in patients that present mutation in BRAF. The combination of BRAFi with MEK inhibitors (MEKi) further improved efficacy and duration of the response. However, most of these patients will develop resistance to BRAFi or BRAFi + MEKi combination therapies within a year (Arozarena and Wellbrock, 2017, Rossi et al., 2019). The mechanisms of resistance are pleiotropic and not fully characterised. In general, they result in the reactivation of ERK signalling, and include the paradoxical activation of wildtype (wt) RAF isoforms, secondary mutations and genetic and epigenetic changes that result in a rewiring of signalling networks (Arozarena and Wellbrock, 2017, Rossi et al., 2019). In addition, reactivation of the PI3K/AKT pathway and upregulation of anti-apoptotic signals also can convey BRAFi resistance (Perna et al., 2015, Serasinghe et al., 2015).

The Hippo/MST2 pathway (hereto MST2 pathway), plays an important role in the regulation of proliferation, organ size and cell death (Fallahi et al., 2016, Pan, 2010). Originally discovered in *Drosophila melanogaster* the pathway has functionally diversified in mammals, where it can drive both cell proliferation and cell death (Romano et al., 2014a, Fallahi et al., 2016). In mammals its core element is a kinase module consisting of MST1/2 which phosphorylates and activates LATS1/2, a kinase with multiple substrates including the YAP transcription regulator, which is a main effector of the canonical Hippo pathway discovered in *Drosophila melanogaster*. However, the mammalian MST1/2 can receive inputs from a variety of upstream cues and transmit signals via various downstream effectors in addition to YAP. The pathway’s involvement in the regulation of apoptosis is stimulated by the FAS death receptor and by RASSF1A (O’Neill et al., 2004, Matallanas et al., 2007, Matallanas et al., 2011b), a tumour suppressor gene, whose expression is frequently silenced in cancer (Richter et al., 2009). These proapoptotic signals are antagonized by RAF1, which binds to and inhibits MST2 (O’Neill et al., 2004). RASSF1A competes with RAF1 for MST2 binding causing the release of MST2 from RAF1, allowing MST2 activation, binding to LATS1/2 and subsequent promotion of apoptosis. Interestingly, apoptosis can proceed through two routes. One is mediated by MST2-LATS1 signalling inducing the formation of a YAP-p73 transcriptional protein complex that promotes the expression of pro-apoptotic genes (Matallanas et al., 2007). The other pathway, also mediated by MST2-LATS1 is independent of YAP and leads to the stabilization of the p53 tumour suppressor protein (Matallanas et al., 2011b).

Different lines of evidence indicate that the deregulation of members of the MST2 pathway plays a role in the development of malignant melanoma. YAP1 has been proposed to behave as an oncogene in melanoma (Thompson, 2020). RASSF1A is commonly lost in melanoma patients due to DNA methylation (Reifenberger et al., 2004). LATS1 levels seem to be decreased through different mechanism that include post-translational modification and long non-coding RNAs (Han et al., 2021, Yuan et al., 2015). These results suggest that loss of a functional MST2 pathway might be associated with melanoma development. However, the possible role of the MST2 pathway in the acquisition of resistance to RAFi has not been studied.

Here, we have investigated the association of MST2 pathway signalling and the rewiring of molecular networks that result in the acquisition of resistance to BRAFi in melanoma cell lines. We show that mutant BRAF^V600E^ binds and inhibits MST2 preventing the activation of MST2-dependent apoptosis. In BRAFi resistant melanoma cells developed in our group, we show that LATS1 and MST2 expression reduced due to ubiquitin ligase-dependent degradation. Treatment of resistant melanoma cells with proteasome inhibitors results in a rescue of MST2 and LATS1 expression and restoration of pro-apoptotic signalling. Finally, results from a small cohort of patients with resistance to BRAFi indicate that MST2 downregulation might be associated with the acquisition of resistance in human melanoma.

## Results

### The MST2 pathway is inhibited by mutant BRAF in melanoma cells and the effect is rescued by BRAF inhibitors

The MST2 kinase is an important regulator of cellular growth and proliferation, and abundant evidence shows that deregulation of the MST2 signalling network is associated with cancer development (Fallahi et al., 2016, Pan, 2010). We have previously shown that MST2 mediates a proapoptotic signal that is inhibited by RAF kinases (O’Neill et al., 2004) and were interested in studying the possible role of the MST2 pathway in the response to RAF inhibitors used in the clinic to treat melanoma. Therefore, we treated a panel of melanoma cell lines that included the 2 main driving mutations in melanoma, BRAF^V600E^ (A375, SK-Mel28 and WM-793 cells) and the NRAS^Q61R^ (Sk-Mel2 cells) point mutations with the BRAF inhibitor PLX4032 (vemurafenib) for 1h or 24h. Using phosphorylation of the activating MST1/2 T180 residue as a read out for MST2 kinase activity, the results showed that PLX4032 caused a rapid and sustained activation of MST1/2 in the three BRAF mutant cell lines, while no effect was observed in SK-Mel2 cells that do not respond to BRAF inhibitors (Fig. 1A). Importantly, we observed that overexpression of wt BRAF and the dimerization defective BRAF^R509H^ mutant (Poulikakos et al., 2010) did not inhibit MST2 activity, while expression of the oncogenic mutants BRAF^V600E^ and BRAF^V600E/R509H^ in in HeLa cells caused a complete inhibition of MST1/2 basal activation (Fig. 1B). These results indicated that MST2 might be regulated by BRAF^V600E^ in melanoma cells, and that this does not require BRAF dimerization. The MST2 network crosstalks with the ERK pathway. MST2 can induce the phosphorylation of RAF1 at the inhibitory S259 site (Romano et al., 2014a), while both RAF1 and oncogenic BRAF can bind to and inhibit the core kinases MST2 and MST1, respectively (Matallanas et al., 2007, O’Neill et al., 2004, Lee et al., 2011). Therefore, we tested if mutant BRAF also binds MST2. Co-immunoprecipitation assays demonstrated that MST2 readily bound to mutant BRAF^V600E^ in melanoma cells while overexpressed wild type BRAF only showed a weak interaction (Fig. 1b). The binding of mutant BRAF^V600E^ was confirmed using a BRAF^V600E^ mutant specific antibody (Fig. 1C). Reversing the IP by immunoprecipitating MST2 also detected strong association of MST2 with BRAF in cell line expressing the oncogenic mutant and no association in cells that express only wt BRAF cells. Interestingly, in all mutant BRAF cell lines we observed a clear decrease of interaction of MST2 and BRAF^V600E^ when the cells were treated with PLX4032. A concomitant increase of LATS1-YAP interaction is shown after 1 hour of treatment with BRAFi (Fig. S1). Taken together, these observations indicate that PLX4032 may activate MST2 pathway by releasing it from BRAF^V600E^ inhibitory binding. If this hypothesis is correct, downregulation of MST2 expression in mutant BRAF cells should prevent PLX4032 induced cell death. Indeed, using siRNA to knock down MST2 by ca. 70% in the three mut BRAF cell lines reduced PLX4032 induced apoptosis between 70-50% (Fig. 1D).

**Figure 1.**
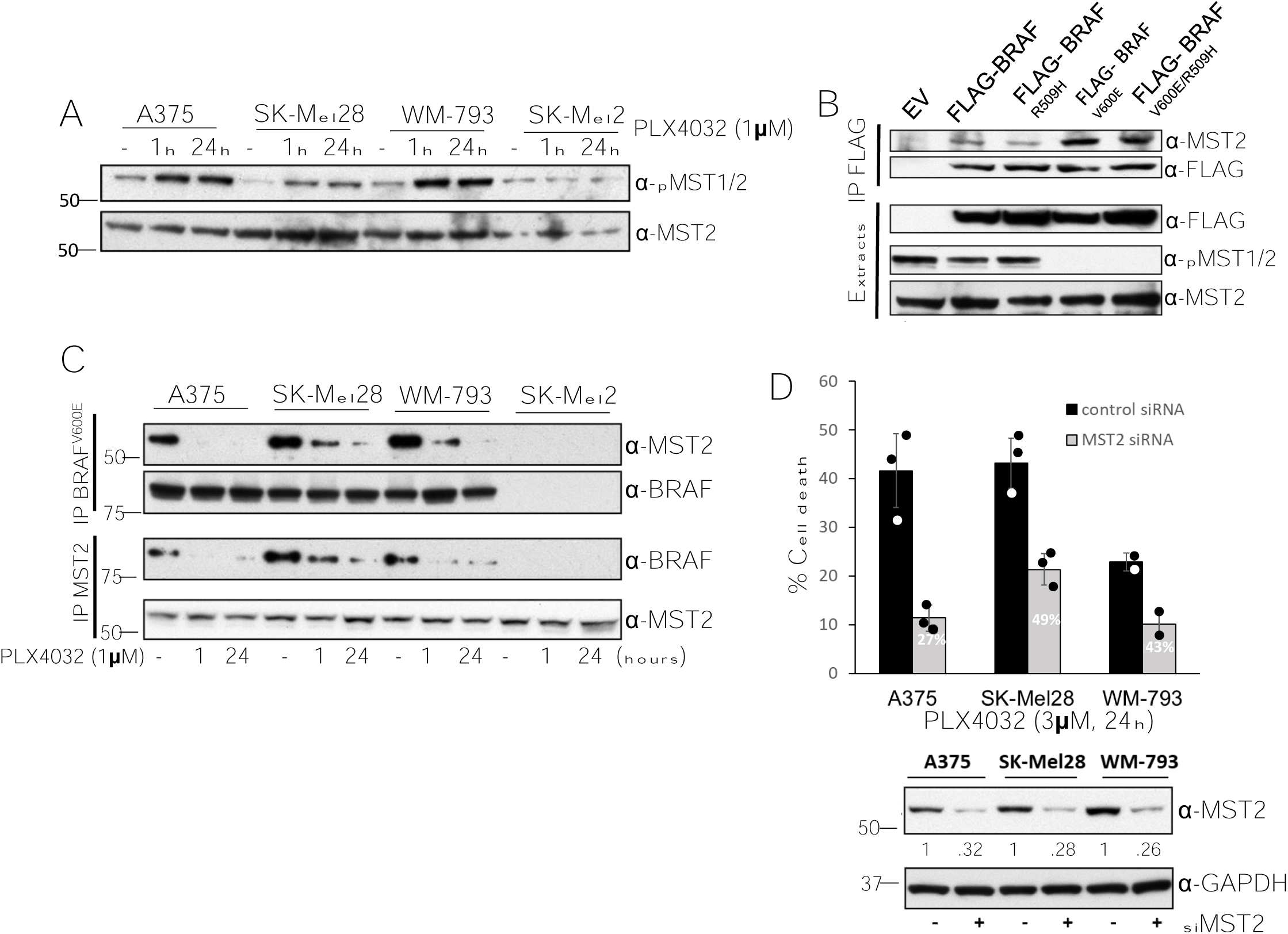
BRAF^V600E^ inhibits MST2 signalling. **A)** A375, SK-Mel28, WM-793 and SK-Mel2 cell were treated with 1 µM PLX4032 for the indicated times. Cell extracts were analysed by immunoblotting with the indicated antibodies. **B)** HeLa Cells were transfected with pCDNA-FLAG (EV) or the different FLAG-BRAF constructs indicated. After 48 hours cells were lysed and FLAG-BRAF protein complexes were immunoprecipitated using FLAG beads. BRAF-MST2 interaction and MST2 activation were determined by immunoblotting with the indicated antibodies. **C)** A375, SK-Mel28, WM-793 and SK-Mel2 were treated with PLX4032 for the indicated times before MST2 and BRAF^V600E^ were immunoprecipitated. Endogenous MST2 and BRAF interaction was monitored by blotting with the indicated antibodies. **D)** Mutant BRAF cells A375, SK-Mel28 and WM-793 were transfected with non-targeting (control) or MST2 siRNA (50 nM). 24 hours after transfection the cells were treated with PLX4032 (3 μM) for 24 hours and cell death was determined by assaying DNA fragmentation by FACS. Percent apoptosis in cells transfected with MST2 siRNA relative to control transfected cells is indicated. An aliquot of the cells was lysed and immunoblotted with the indicated antibodies to determine protein expression. The expression of MST2 normalised to the GAPDH loading control was determined by Image J scanning. Error bars show standard deviation n=3 for A375 and SK-Mel28; n=2 WM-793.

These results show that mutant BRAF interacts with MST2 and inhibits its proapoptotic signal. Moreover, the results suggest that breaking up the interaction between mutant BRAF and MST2 causing MST2 activation is part of the mechanism of action of PLX4032.

### Acquisition of resistance to BRAF inhibitors causes the downregulation of MST2 pathway proteins

Most melanoma patients develop BRAF inhibitor resistance within a year after initiation of treatment. This is due to the paradoxical activation of RAF isoforms or a rewiring of the signalling network that circumvents the RAF blockade to achieve ERK activation (Matallanas et al., 2011a, Arozarena and Wellbrock, 2019). In light of the observation that the MST2 pathway participates in mediating cell death in PLX4032 sensitive mutant BRAF melanoma cells, we wanted to know if this pathway is associated with the acquisition of resistance to BRAF inhibitors. To study this, we generated resistant cell lines from the A375, SK-Mel20 and WM-793 cells by growing the cells in media containing increasing amounts of PLX4030 for a period of 6 months (Fig. 2A). Cells that could sustain proliferation in the presence of 3 µM PLX4032 were considered resistant. We obtained several PLX4032 resistant clones (Figure 2A and S2A). The resistant clones showed lower level of activation of ERK than the parental cells but, importantly while treatment with PLX4030 causes a complete inhibition of ERK in parental cells, the resistant clones could maintain ERK phosphorylation (Fig. 2A). These results are similar to other reports of BRAF inhibitor resistant cells, confirming that the acquisition of resistance is due to a re-activation of ERK signalling (Lee et al., 2020). The resistant cells also had lost sensitivity to PLX4032 induced apoptosis (Fig. 2B). As all clones showed similar responses in terms of ERK re-activation and resistance to apoptosis, we selected one representative clone from each resistant cell line to perform most of the follow up experiments. Importantly, MST2 expression levels were downregulated in all the resistant clones suggesting that losing the pro-apoptotic function of MST2 signalling is involved in the acquisition of resistance to BRAF inhibitors (Fig. 2C and Fig. S2A, B). Therefore, we assessed the protein expression of LAST1, RASSF1A and YAP1 in the resistant cells, observing a strong decrease in their expression in resistant cells (Fig. 2C). The only exception was YAP1 expression in WM-793 cells, which did not change in the resistant cells. A375 and SK-Mel28 cells do not express RASSF1A, but RASSF1A is expressed in WM-793 cells and reduced in their resistant variants. As expected from the reduction of protein abundances, the basal levels of MST2 and LATS1 activation were reduced in the resistant cells. Importantly, treatment with PLX4032 could not activate these kinases as it did in the parental cells. These data clearly indicate that both the abundance and activation of MST2 pathway proteins are downregulated during the acquisition of RAF inhibitor resistance.

**Figure 2.**
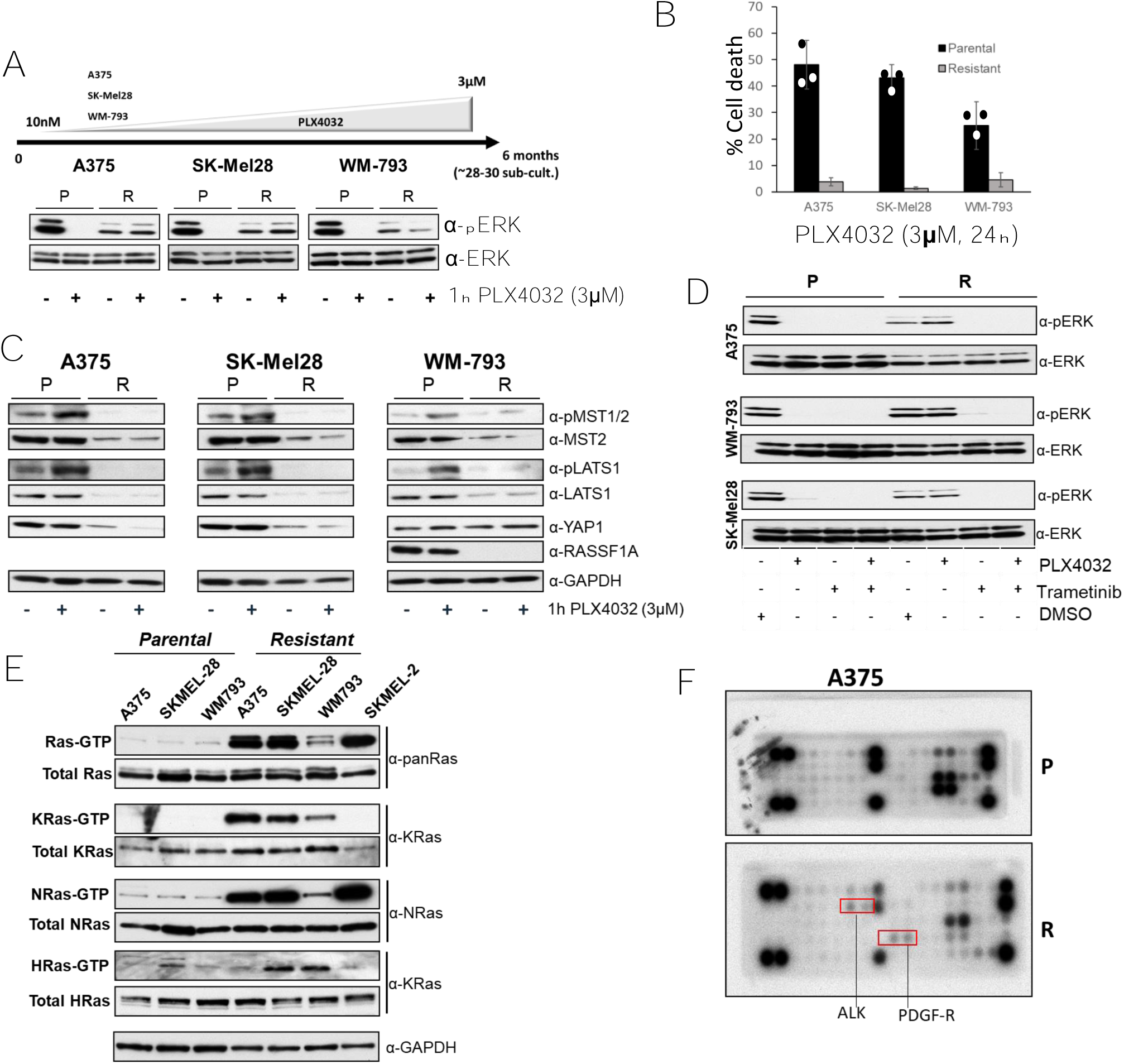
Mutant BRAF melanoma cells resistant to PLX4032 cells rewire ERK and MST2 pathways. **A)** A375, SK-Mel28 and WM-793 cell were grown in increasing concentration of PLX4032 for 6 months. Parental (P) and Resistant (R) cells grown without PLX4032 for 48 hours were treated with PLX4032 (3 µM) for 1 hour and ERK activation was determined by immunoblotting with the indicated antibodies. **B)** Parental and resistant A375, SK-Mel28 and WM-793 cells were treated with PLX4032 (3 µM) for 24 hours and DNA fragmentation was determined but PI staining using FACS. Error bars show SD, (n=3). **C**) Parental and resistant cells were treated as in A) and the expression of MST2 pathway proteins and GAPDH as loading control were determined by immunoblotting with the indicated antibodies. **D)** Parental and PLX4032 resistant cells were treated with PLX4032 (3 μM) and/or trametinib (10 nM) for 1 hour. Cell extracts were immunoblotted with the indicated antibodies. **E)** Lysates from serum deprived (16 hours) parental and resistant cells were incubated with GST-RBD to determine the amount of RAS GTP. A lysate aliquot was used to determine total RAS expression. Expression of RAS isoforms was determined by immunoblotting with the indicated antibodies. **F)** Parental (P) and resistant A375 (R) cells were lysed and cell extracts were incubated with PathScan® RTK Signaling Antibody Arrays. Red boxes show ALK and PDGF-receptor spots.

To further characterise the rewiring mechanisms occurring during the development of resistance to PLX4032 in our cell lines we used BRAF and MEK inhibitors to probe the activation of the ERK pathway. All resistant cells show an inhibition of ERK1/2 phosphorylation when the cells are treated with MEK inhibitor indicating that there was not secondary mutation of MEK in the resistant cell lines (Fig. 2D). By contrast, PLX4032 failed to block ERK activation, presumably due to the induction of BRAF-RAF1 heterodimers (Fig. S2C), which have been shown to maintain ERK signalling or even cause paradoxical pathway activation (Arozarena and Wellbrock, 2017, Rossi et al., 2019). PLX4032 combined with MEK inhibition caused ERK inhibition (Fig. S2C) in keeping with the synergy observed between RAF and MEK inhibitors (Sturm et al., 2010, Flaherty et al., 2012). Enhanced RAF dimerization is a main source of resistance to BRAFi in melanoma and is frequently caused by RAS activation (Johnson et al., 2015). Indeed, RAS activation was clearly increased in all three resistant cell lines, especially in A375 and SK-Mel28 resistant cells which showed similar levels of RAS activation as the mutant NRAS cell line SK-Mel-2 (Fig. 2E). Using isoform specific antibodies, we observed that A375 cells featured a strong activation of KRAS and NRAS but not HRAS. In SK-Mel28 resistant cells all RAS isoforms were hyperactivated with respect to the parental cells. Finally, in WM-793-R cells there was a weaker activation of NRAS compared to the other resistant cell lines, but KRAS and HRAS were clearly hyperactivated compared to the parental cell lines. This increase of RAS signalling could be caused by secondary mutations in RAS isoforms or more likely by the activation of upstream regulators, which causes hyperactivation of several RAS isoforms. To test this, we used PathScan receptor tyrosine kinase (RTK) signalling antibody arrays. All cell lines had upregulated RTK signalling. A375R showed an increase of PDGF-R and ALK. SK-MEL28 showed an increase IRS1 and SRC activation (Fig. 2F and S2D). Finally, WM793R had an increase of IGF-1R and the SRC effector SRC. These data indicate that the increase of activation of RAS isoforms may be caused by the activation of upstream receptor signalling in the resistant cells. The fact that the three cell lines showed activation of different receptors may explain why there is a differential activation of the RAS isoforms since we and other have shown that the isoforms are differentially regulated by upstream signals (Kiel et al., 2021).

In summary, the resistant cell lines that we have generated have acquired significant changes in the signalling machinery including diverse changes in RTK and RAS signalling, but similar changes in the ERK and MST2 pathways.

### MST2 pathway proteins are downregulated by proteasome degradation in BRAFi resistant cells

These results indicated that a decrease of expression of MST2 pathway proteins is involved in BRAFi resistance. Therefore, we tried to identify the mechanisms behind this downregulation in the BRAF^V600E^, PLX4032 resistant cell lines.

Loss of expression of RASSF1A in A375 is due to DNA methylation of the gene promotor (Yi et al., 2011), and the expression of MST1/2 and LATS1/2 in cancer also can be regulated by DNA methylation (Fallahi et al., 2016). Therefore, we first tested whether the expression changes of core Hippo proteins were caused by decreased gene transcription. Measuring mRNA expression of MST1/2, LATS1/2 and YAP1 by RT-PCR showed equal or upregulated expression in all resistant cell lines (Fig. 3A). This makes it unlikely that the reduction of protein expression is due to gene silencing or a reduction in gene expression.

**Figure 3.**
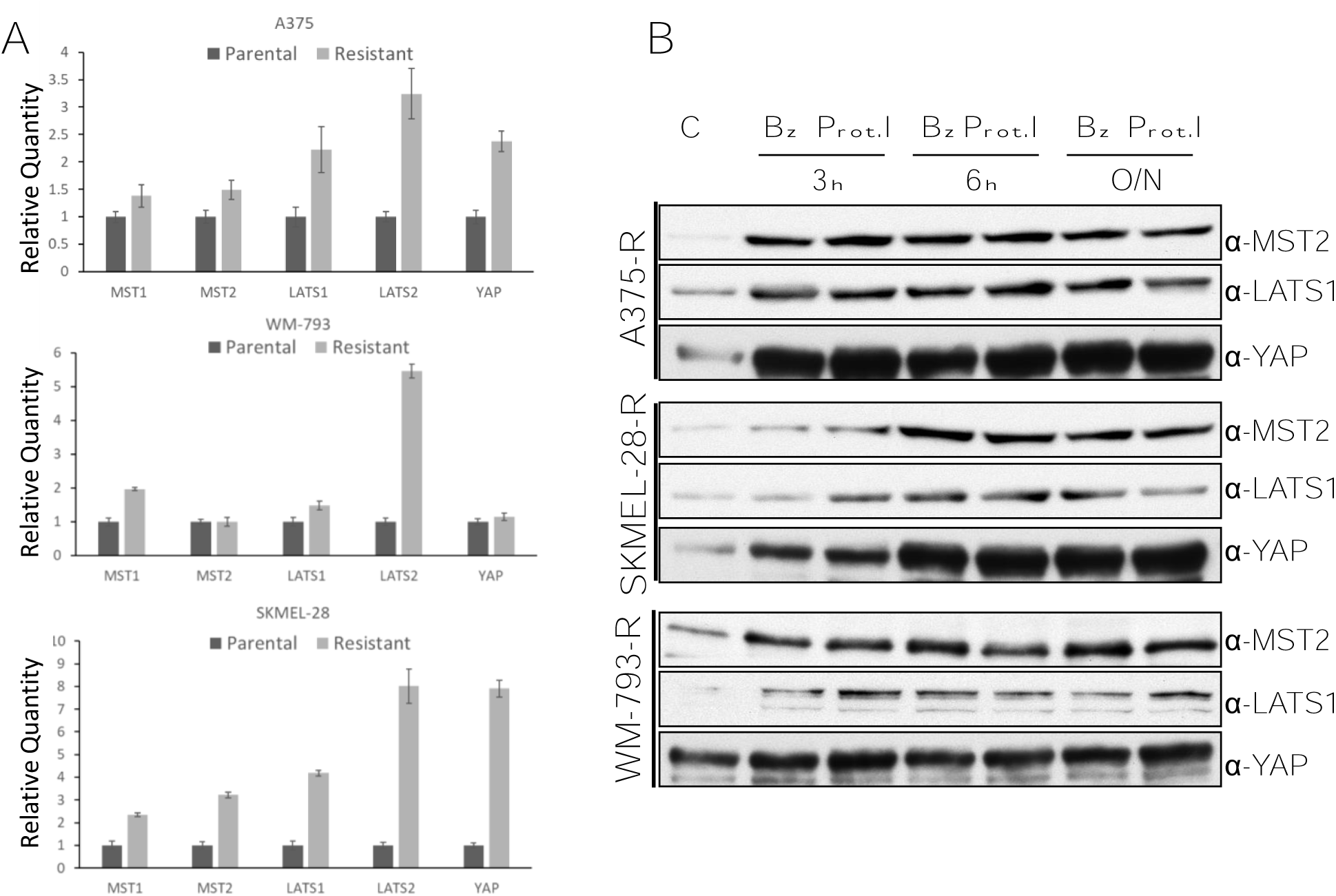
MST2 pathway proteins are downregulated by proteasomal degradation in PLX932 resistant cells. **A)** RNA was extracted from parental and resistant A375, SK-Mel28 and WM-793 cells and the level of expression of the indicated RNAs was determined by qRT-PCR. Error bars show SD for 3 measurements. **B)** Resistant A375, SK-Mel28 and WM-793 cells (C) were treated with Bortezomib (Bz, 50 nM) or Proteasome inhibitor I (Prot.I, 10 µM) for the indicated times (O/N, overnight). Cell lysates were blotted with the indicated antibodies.

Overexpression of a mRNA can be a compensatory mechanism to counterbalance the enhanced degradation of the cognate protein (Prelich, 2012). To test this possibility, we treated the cells with the proteasome inhibitors bortezomib and Proteasome inhibitor 1. Both inhibitors consistently and rapidly increased the expression of MST2, LATS1 and YAP1 in all three resistant cell lines (Fig. 3B). These data indicated that the reduction in MST2, LATS1, and YAP1 proteins expression in PLX4032 resistant cells is due to the acquisition of resistance to the BRAF inhibitors the core proteins of the Hippo pathway are targeted for proteasomal degradation.

### Increased ubiquitination promotes MST2 and LATS1 proteasomal degradation in BRAFi resistant cell lines

Proteasomal degradation usually is preceded by ubiquitination, which directs the target protein to the proteasome (Kornitzer and Ciechanover, 2000). To test if there was an increase of ubiquitination of MST2 and LATS1 in the PLX4032 resistant cells, we co-transfected the cells with HA-tagged ubiquitin and FLAG-tagged MST2 and determined changes of ubiquitination by immunoprecipitating FLAG-MST2. Despite similar ubiquitination levels in the total cell lysates, MST2 ubiquitination was clearly enhanced in the resistant cell lines and correlated with lower expression levels of FLAG-MST2 in the cell extracts (Fig. 4A). Similarly, expression of Myc-tagged LATS1 in WM793 and SK-Mel 28 cells showed increased ubiquitination in the resistant variants (Fig. S3). These experiments suggest that the expression of MST2 and LATS1 in PLX4032 resistant cells is downregulated by ubiquitin-dependent proteasomal degradation.

**Figure 4.**
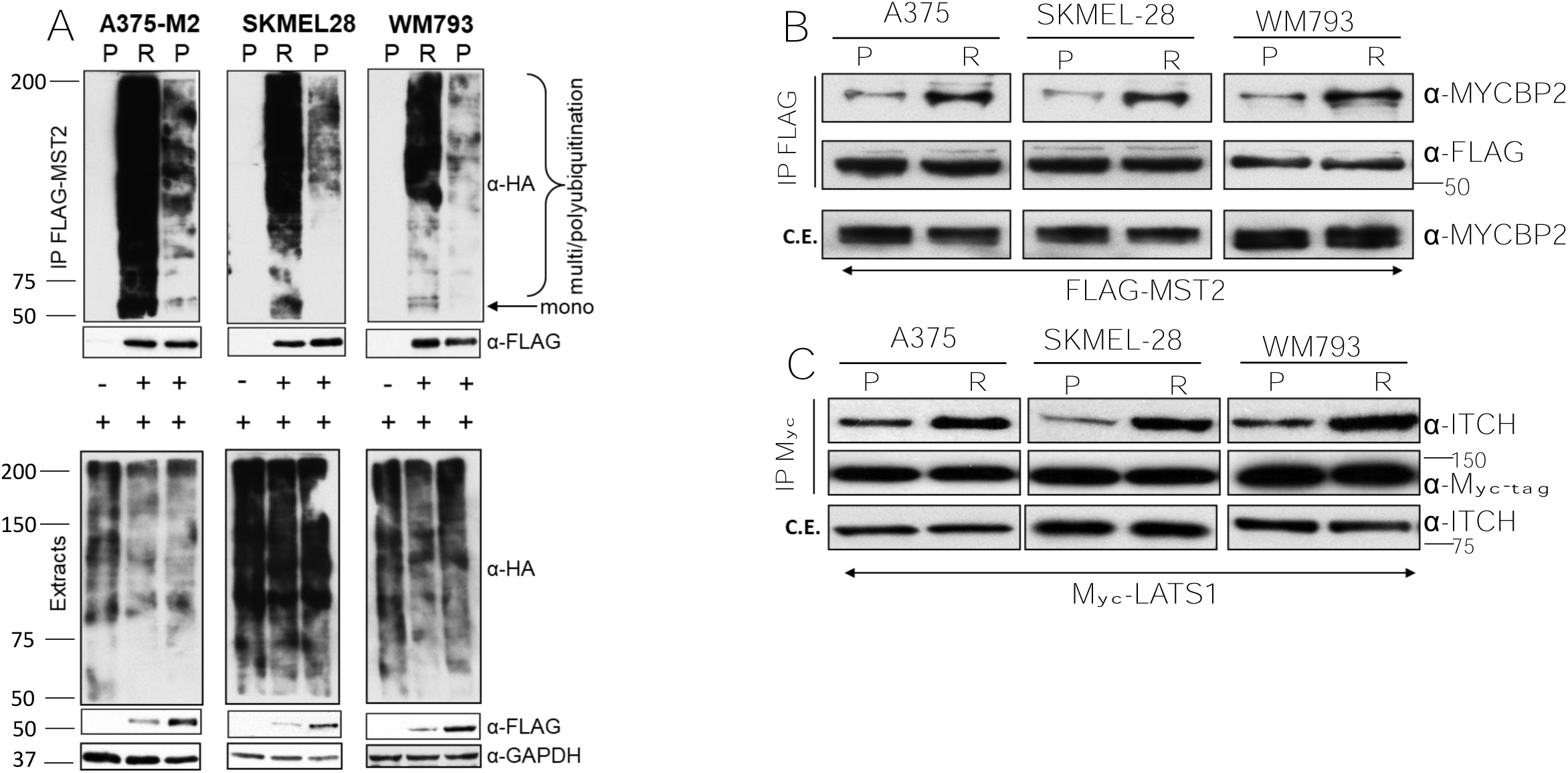
MST2 and LATS1 exhibit increased ubiquitination and association with E3-ubiqitin ligases in resistant melanoma cells. **A)** Parental and resistant A375, SK-Mel28 and WM-793 cell lines were transfected with HA-ubiquitin (1 μg) alone or together with FLAG-MST2 (0.5 µg) and lysed 48 hours after transfection. Cell extracts were immunoprecipitated with FLAG antibody beads and ubiquitination of precipitated MST2 was determined using HA antibody. Expression of proteins was determined in cells extract by immunoblotting with specific antibodies. **B)** Parental and resistant cells were transfected with FLAG-MST2 (0.5 µg) and exogenous protein was IP 48 hour after transfection using FLAG beads. Interaction with endogenous MCYBP2 was determined by immunoblotting with specific antibody. **C)** Parental and resistant cells were transfected with Myc-LATS1 (1 µg) and cell lysates were incubated with Myc-tag antibody to immunoprecipitate Myc-LATS1. ITCH interaction with Myc-LATS1 was determined using a specific antibody.

In an attempt to characterise the E3 protein ligases responsible for this ubiquitination we mined two interaction proteomic datasets produced in our group that have been previously used to identify interactors of these kinases (Quinn et al., 2021, Novacek et al., 2020). In a FLAG-MST2 AP-MS experiment we identified MYCBP2 as a binding interactor of MST2. MYCBP2 is an atypical E3 Ubiquitin-Ligase that catalyses the ubiquitination of Threonine and Serine rather than Lysine residues (Pao et al., 2018). Interestingly, when validating the MST2-MYCBP2 interaction by co-immunoprecipitation – Western blotting (coIP-WB), we not only observed the interaction in the parental cell lines, but also saw a clear increase of this interaction in the resistant cells (Fig. 4B). Similarly, we identified the E3 ubiquitin ligase ITCH as a LATS1 interactor in FLAG-LATS1 IPs (Fig. 4C). This interaction was also enhanced in resistant cells. These data indicate that these E3-Ligases might be responsible for the increase of MST2 and LATS1 ubiquitination in the resistant cells. The mechanism seems to be enhanced binding, as the total expression levels of MYCBP2 and ITCH were unchanged between parental and resistant cells.

### Proteasome inhibition promotes cell death in MM resistant cell lines though and MST-dependent mechanism

The previous data suggested that ubiquitination and proteasomal degradation of MST2 pathway proteins play an important role in the acquisition of resistance to BRAFi. This was further supported by the observation that the proteasome inhibitor bortezomib selectively induced PARP cleavage (a marker of apoptosis) in BRAFi resistant A375-R and SKMEL28-R cells but not in parental cells (Fig. 5A). In WM793 cells PARP was overexpressed in resistant cells and showed constitutive cleavage which was increased by bortezomib. Based on this observation we hypothesised that the enhanced degradation of MST2 pathway proteins prevents BRAFi induction of apoptosis in resistant cells, and that bortezomib can restore apoptosis by stabilizing MST2 pathway proteins. In order to test this, we treated the resistant cell lines with increasing amounts of bortezomib. We observed that bortezomib caused a clear dose dependent increase in MST2 protein abundance in all 3 resistant cell lines (Fig. 5B). This was accompanied by an increase in cell death when the cells were treated with more than 30nM of bortezomib (Fig. 5C). Expression of specific MST2 siRNA significantly reduced bortezomib induced cell death further confirming the role of the MST2 pathway in mediating the response to bortezomib treatment. Of note, we also tested whether bortezomib can re-sensitize resistant cells to PLX4032, but the combination was toxic with a very small therapeutic window making a clear assessment of possible re-sensitisation or synergy difficult.

**Figure 5.**
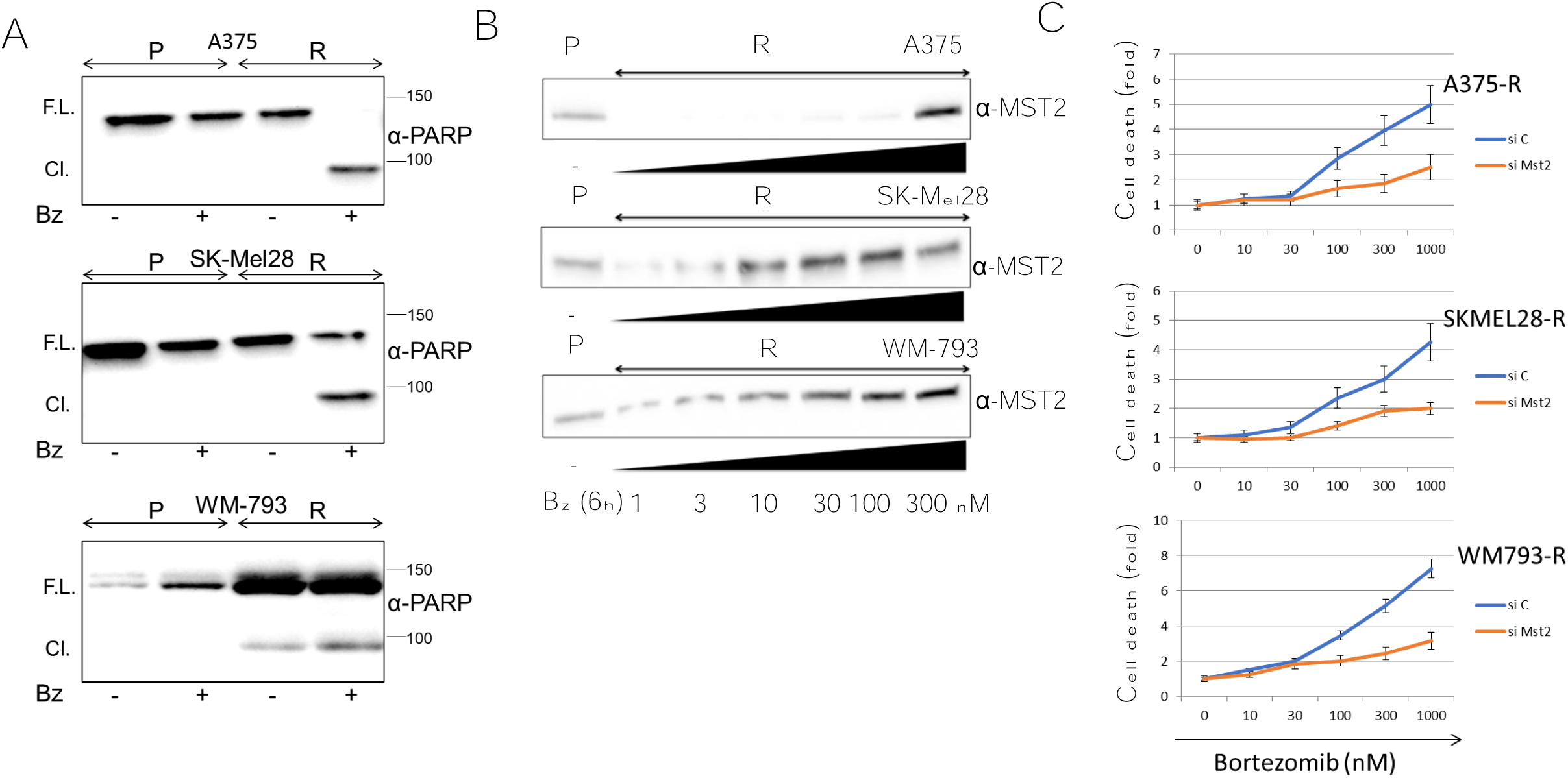
Bortezomib induce apoptosis in PLX4032 resistant cells in an MST2 dependent fashion. **A)** Parental and resistant A375, SK-MEL28 and WM-793 were treater with bortezomib (10 nM) for 6 hours and cell lysates were immunoblotted with PARP antibody to determine caspase expression and cleavage. **B)** A375, SK-MEL28 and WM-793 resistant cell lines were treated with the indicated concentrations of bortezomib and MST2 expression levels were monitored by immunoblotting with specific antibody. **C)** A375, SK-MEL28 and WM-793 resistant cell lines were transfected with non-targeted (si C) or MST2 siRNA (siMST2) and after 24 hour of transfection the cells were treated with the indicated concentrations of bortezomib for 3 hours and cell death was determined by assaying DNA fragmentation by FACS. Error bars show SD, n=3.

### Loss of expression of MST2 may correlate with development of resistance to BRAFi in patients

The data described above indicate that the reduction of MST2 pathway protein expression contributes to the development of resistance to vemurafenib. As a first attempt to test if this might be relevant in the clinical setting, we used a small cohort of nine human metastatic melanoma patients who were resistant to vemurafenib treatment alone or in combination with MEK inhibitors, and where both pre-treatment and relapse histological samples were available from previous studies (Frederick et al., 2013) (Fig. 6, S4, S5 and Table 1,). While MST2 expression was detectable in melanomas before treatment, it was lost or reduced in 8 out the 9 patients, who relapsed or progressed on vemurafenib treatment. Although the sample size is small, these data suggest that a reduction in MST2 expression is part of the mechanism how melanomas develop resistance to BRAFi in patients.

**Table 1.**
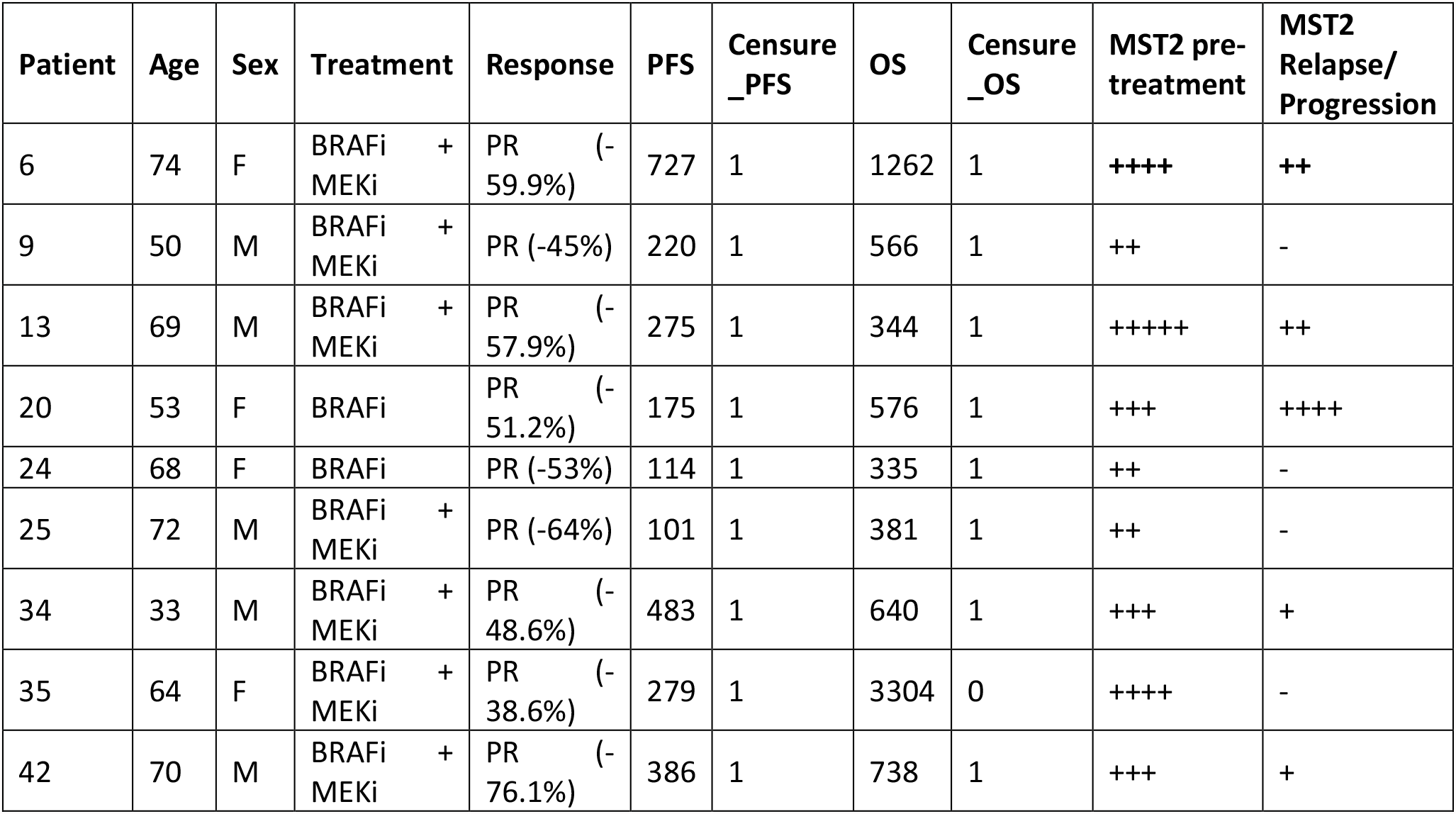
Loss of MST2 in 8 out of 9 patients with relapsed or progressive disease. Histopathological slides of tumour samples taken before treatment and upon disease relapse or progression were stained for MST2 expression. MST2 expression is indicated by “+” on a scale from 1-5 (weak – very strong); “-” means no MST2 expression detectable. PFS progression free survival; OS overall survival. PFS and OS indicate days relative to treatment start date. Censure_PFS: 1=progressed, 0=not progressed. Censure_OS=dead, 0=alive. PR partial response. BRAFi treatment with vemurafenib. MEKi treatment with tramatenib and dabrafenib.

**Figure 6.**
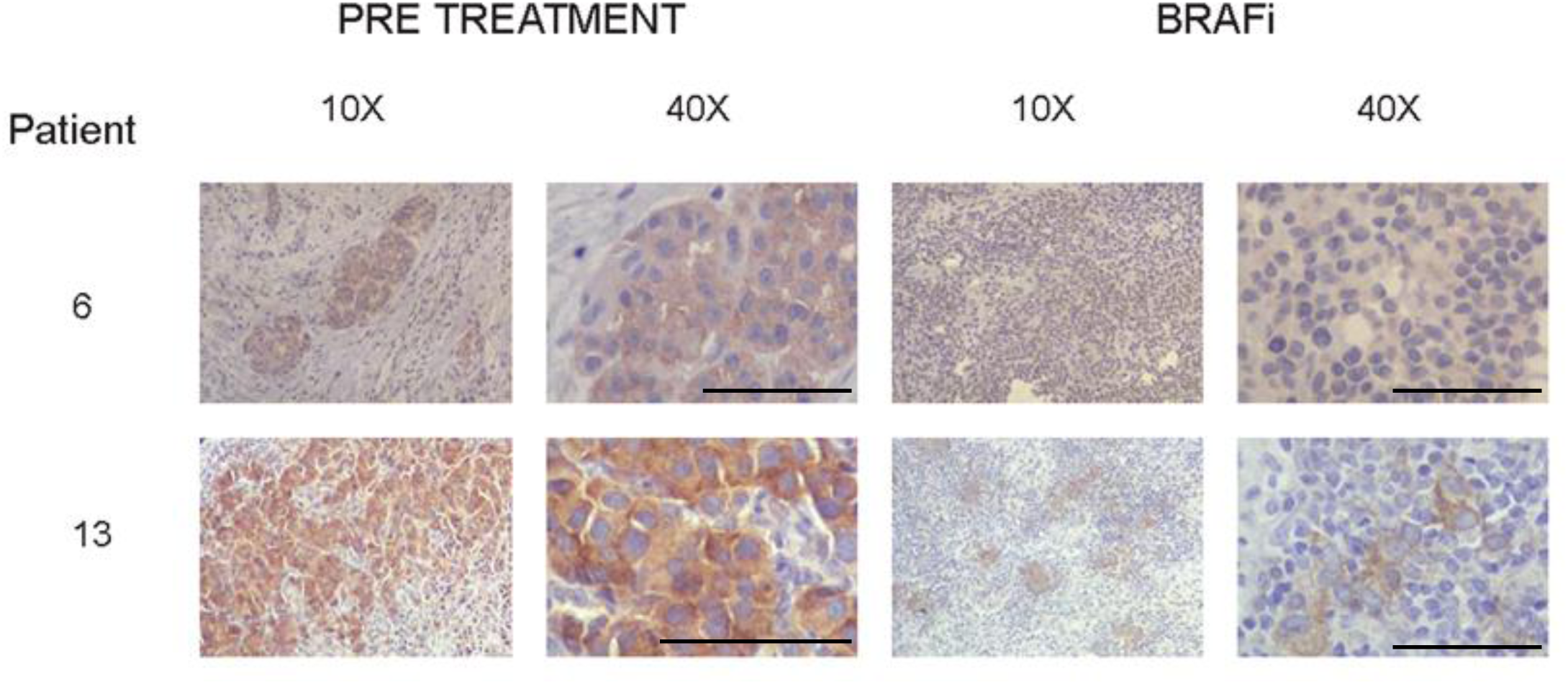
MST2 expression is decreased in patients with metastatic melanoma treated with BRAF inhibitors. Immunohistochemistry (10x and 40X magnification) for the MST2 expression was detected as explained in materials and methods. Tumours biopsied from representative patients were stained with haematoxylin and eosin. The figure shows expression of MST2 in patients 6 and 13 before treatment with BRAF inhibitor (pre-treatment) and after treatment with BRAF and MEK inhibitors (BRAFi). Bar=100 μm

## Discussion

Malignant cutaneous melanoma is hallmarked by a hyperactivation of the RAS-RAF-MEK-ERK pathway that drives several aspects of malignant transformation, including proliferation, survival, invasiveness and metastatic spread. All three most common driver mutations (BRAF, NRAS and NF1 mutations) converge on activating ERK signalling (Conway et al., 2020). About 10 years ago the introduction of vemurafenib revolutionized the treatment of metastatic melanoma patients with BRAF^V600E^ mutations (Bollag et al., 2010). However, despite good response rates and remarkable efficacy, most patients became resistant within 6-8 months. The addition of MEK inhibitors synergizes with BRAFi (Sturm et al., 2010) improving the clinical efficacy and delaying the onset of resistance by ca. 3 months (Flaherty et al., 2012). Later BRAFi generations suffered from the same problem, which triggered numerous studies analysing resistance mechanisms (McKenna and García-Gutiérrez, 2021, Lee et al., 2020, Czarnecka et al., 2020, Proietti et al., 2020, Rossi et al., 2019, Sanchez et al., 2018). Despite a variety of mechanisms were discovered the large majority shares the re-activation of ERK signalling as common theme. However, another interesting observation was that the various ways to reactivate ERK rested mainly on network rewiring rather than mutations that prevented drug binding to the target. This finding prompted us to investigate the role of pathways that crosstalk with RAF.

While phosphorylation and activation of MEK1/2 is the only widely accepted catalytic function of the RAF kinases, they can regulate other signalling pathways independent of their kinase activity (Rauch et al., 2011, Baccarini, 2005, Nolan et al., 2021). These include the suppression of proapoptotic pathways mediated by ASK1 (Chen et al., 2001) or MST2 (O’Neill et al., 2004). The connection the MST2 pathway seemed particularly interesting, because several components of the MST2 pathway have been previously implicated in melanoma albeit in different roles. Overexpression of YAP1, a main transcriptional effector of MST2 signalling, can drive melanomagenesis (Thompson, 2020). By contrast, RASSF1A, an upstream activator of MST2, and LATS1, a downstream substrate of MST2, may work as tumour suppressors. RASSF1A expression is commonly lost in melanoma patients (Reifenberger et al., 2004), and the expression of LATS1 is often downregulated (Han et al., 2021, Yuan et al., 2015). These observations are consistent with a tumour suppressive function of MST2 signalling that is independent of YAP1. Interestingly, we have previously identified such a pathway where MST2-LATS1 signalling leads to the stabilization of the TP53 tumour suppressor protein and subsequent apoptosis in a YAP1 independent fashion (Matallanas et al., 2011b). TP53 is mutated in 19% of melanoma patients in the TCGA database, and these mutations seem to be mutually exclusive with MST1/2 amplifications occurring in 7% of melanoma patients although more data is necessary to get statistical confirmation for this inverse correlation. This potentially intriguing relationship warrants a study in its own right.

In the current study we have focussed on the crosstalk between the RAF and MST2 pathways. In our original study (O’Neill et al., 2004) we found that RAF1 could bind to and inhibit MST2, while BRAF was rather inefficient in binding MST2. Here, we confirm the weak binding of BRAF to MST2 but find that BRAF^V600E^ binds strongly to MST2. Vemurafenib treatment releases this inhibitory interaction resulting in MST2 activation and apoptosis that at least in part is MST2 dependent. An inhibitory interaction between BRAF^V600E^ and the related MST1 kinase was observed previously in thyroid cancer (Lee et al., 2011). These results suggest that BRAF^V600E^ has an altered conformation that not only enhances its kinase activity towards MEK but also its ability to inhibit MST2 pro-apoptotic signalling. This may thwart the switch-like relationship between the RAF1 and MST2 pathways, where RAF1 can either activate MEK to drive proliferation or inhibit MST2 to block apoptosis (Romano et al., 2014b). In order to phosphorylate MEK, RAF1 must release MST2 thereby coupling MEK driven proliferation to a higher risk of apoptosis mediated by MST2. As PLX4032 releases MST2 from BRAF^V600E^ leading to apoptosis, it is very plausible that the acquisition of drug resistance not only requires re-activation of ERK signalling but also neutralization of the pro-apoptotic signals mediated by the MST2 pathways. Our results suggest that PLX4032 resistant cells can solve this dilemma by downregulating the expression of MST2 and LATS1 proteins. This seems not only happening in cell lines but also in melanoma patients emphasizing the urgency to identify a mechanism and potential target for therapeutic interference.

Our results suggest that MST2 and LATS1 protein expression in resistant cells is downregulated due to enhanced ubiquitination and subsequent proteasomal degradation. Using AP-MS we also identified the ubiquitin ligases as MYCBP2 and ITCH, respectively. MYCBP2 is an unconventional E3 ubiquitin ligase that modifies threonine and serine instead of lysine residues (Pao et al., 2018). It has few known substrates, suggesting a high substrate specificity, and an unusual catalytic mechanism (Pao et al., 2018), which may make it an attractive drug target. Intriguingly, MYCBP2 is highly expressed in skin (Uhlén et al., 2015), and the phosphorylation of the major MYCBP3 phosphorylation site, serine 3503, is downregulated by both PLX4032 as well as the MEK inhibitor AZD6244 (selumetinib) (Stuart et al., 2015). Unfortunately, the functional consequence of this phosphorylation is unknown, but suggests that the RAF pathway may regulate MYCBP2 and thereby influence MST2 degradation. ITCH is a classic E3 ubiquitin ligase that can ubiquitinate RASSF1A (Pefani et al., 2016) and LATS1 (Ho et al., 2011, Salah et al., 2011) and mark them for degradation. The ITCH mediated reduction of RASSF1A and/or LATS1 expression promotes cell proliferation, epithelial-mesenchymal transition, and tumorigenicity. Vice versa, knocking down ITCH expression increased LATS1 expression resulting in reduced proliferation and increased apoptosis (Salah et al., 2011, Ho et al., 2011). Thus, ubiquitination plays an important role in regulating the biological activity of the MST2 pathway. Importantly, the relevance of ubiquitin degradation of the kinases of the pathway is supported by a recent study showing that the E3 Ubiquitin ligase RNF6 interact with MST1 promoting the degradation of this kinase in breast cancer which is associated with short survival (Huang et al., 2022).

Interestingly, the total abundance of ITCH and MYCBP2 was not altered in resistant cells, suggesting that the enhanced association between them and MST2 and LATS1 may depend on affinity changes caused by posttranslational modifications (PTMs). This will require mapping the exact binding sites and PTMs that map to these sites or can alter their conformation. These analyses may reveal new druggable targets that could be used to overcome BRAFi resistance in melanoma. Here, we have concentrated on restoring MST2 and LATS1 expression using proteasome inhibitors. Indeed, elevation of MST2 and LATS1 protein concentrations induced apoptosis in PLX4032 resistant melanoma cells. Combination with PLX4032 proved very toxic preventing us to determine whether these combinations are synergistic (data not shown). This is in contrast to thyroid cancer, where PLX4032 and Bortezomib synergised to inhibit BRAF^V600E^ transformation (Tsumagari et al., 2018). Thus, in melanoma rather than a restoration of sensitivity to PLX4032, proteasome inhibition and elevation of MST2 and LATS1 protein abundances seems sufficient for triggering apoptosis. Interestingly, pharmacological screens have pointed to proteasome inhibitors selectively being active against cancer cells expressing BRAF^V600E^ (Zecchin et al., 2013). In summary, our findings show that the crosstalk between the RAF and MST2 pathways could provide a target that may overcome resistance to BRAFi.

## Materials and methods

### Cell culture and transfection

A375, SK-Mel28, WM-793 and SK-Mel2 were obtained from ATCC and were validated by sequencing before initiation of the study. All cells were grown in RPMI media (Gibco, USA) supplemented with 10% foetal bovine serum (Gibco) and 2 mM L-glutamine (Gibco). Media for the resistant cells contains 2 μM PLX4032 (Selleck chemicals, USA). HeLa cells were also validated by sequencing and grown in DMEM (Gibco) 10% foetal bovine serum and 2 mM L-glutamine. The cells were maintained at 37°C and 5% CO_2_. Cells were transfected with Lipofectamine 2000 (Invitrogen, USA) following manufacturers protocol and the amount of DNA and siRNA is indicated in each experiment. FLAG-BRAF, -BRAF^R509H^, -BRAF ^V600E/R509H^, and BRAF^V600E^, MYC-LATS1 and FLAG-MST2 have been described (Romano et al., 2014a, Quinn et al., 2021, Jambrina et al., 2016) HA-Ubiquitin was a gift from Edward Yeh (Addgene plasmid # 18712)(Kamitani et al., 1997). MST2 SMARTpool siRNA is from Dharmacon (M-012200, CO, USA) and was validated before (Matallanas et al., 2007). Bortezomib (S1013), trametinib (S2673) are from Selleck Chemicals and Proteasome inhibitor I from Calbiochem (539160, CA, USA).

### Generation of resistant cell lines

A375, SK-Mel28 and WM-793 cells were grown in media containing increasing amounts of PLX4032 for 6 months. The cells were initially grown in 10 nM PLX4032 containing media for 1 month and cells that proliferated in this condition were expanded after 2 weeks subculture 3 times. Subsequently, the cells were grown in 30nM PLX4032 and the same sequence was followed (grow for 3 weeks and subculture 3 times). Cells that resisted these conditions were further expanded following the same culture process in increasing amounts on PLX4032 following this sequence 100 nM, 30nM, 300 mM and 1 μM. Finally, after approximately 6 months (28-30 passes) cells were resistant to 3 μM PLX4032. Resistant clones were split into different plates, frozen and stored liquid nitrogen to create cell stocks.

### Cell lysis, immunoprecipitation, and Western blotting

Cellular extracts from the different cell lines were produced using cell lysis buffer containing 150 mM NaCl (Sigma-Aldrich, MO, USA), 20 mM HEPES (Sigma-Aldrich) pH 7.5, 1% NP-40 NP 40 and proteases inhibitors. Immunoprecipitation was performed by adding primary antibody and 5 µl of agarose beads to cell extracts. 5 µl of FLAG beads (Sigma-Aldrich) were added for FLAG IPs. The mix was incubated rotating for 2 hours at 4°C followed by washes with washing buffer (150 mM NaCl, 20 mM HEPES pH 7.5, 0.5% NP-40). Immunoprecipitates or total cell lysates were separated by SDS-polyacrylamide gel electrophoresis, followed by transfer to Polyvinylidene Fluoride (PVDF) membranes for Western blotting. The following antibodies were used: FLAG-M2 (A8592), ERK1/2 (M5670) and phospho-ERK1/2 (M8159), BRAF^V600E^ (SAB5600047) from Sigma-Aldrich; pan-Ras (op40) from Calbiochem; HA-HRP (3F10) from Roche (Germany); HA (sc-7392), MST2 (SC-6213), anti-LATS1 (sc-9388 and sc-12494), YAP1 (SC-15407), BRAF (Sc-5284), HRAS (SC-520), KRAS (SC-30) and NRAS (SC519) from Santa Cruz (CA, USA); anti-RASSF1A (14-6888-82) from ebioscience (CA, USA); STK3 (MST2, ab52641) and MYCBP2 (Ab86078) from Abcam (UK); phospho-MST1/2 (3682), phospho-LATS1 (8654), GAPDH (2118) and Myc-TAG (2276) from Cell Signaling (MA, USA); C-Raf (610152) PARP (556362) and ITCH (611199) from BD transduction laboratories; and ARAF (R14320) from Transduction Lab (NJ, USA).

### Cell death and apoptosis assays

Cell death was analysed by flow cytometry as descried before (O’Neill et al., 2004). Briefly, cells were grown as indicated, and media containing floating cells was collected prior to trypsinization of plates. Trypsinized cells and the collected media were pooled, and cells were pelleted by centrifugation. The cells were washed by resuspension in PBS, and after centrifugation they were fixed with 90% EtOH/PBS for 1 hour. The cells were incubated with propidium iodide and RNAse dissolved in PBS, and the population containing fragmented sub-G1 DNA content was measured using an Accuri C6 Flow Cytometer. In addition, activation of apoptosis was monitored by assessing PARP cleavage using Western blots.

### RTK signalling antibody array

Lysates from parental and resistant cells were used to screen activation of receptor tyrosine kinases using the PathScan® RTK Signaling Antibody Array Kit (7982, Cell Signaling) following the manufacturer’s instructions.

### RAS activity assays

Ras activation was determined by performing RAF-RAS binding Domain (RBD, which binds activated Ras) pulldown assays as previously described (Herrero et al., 2017). Briefly, cells were serum deprived for 16 hours and lysed using magnesium rich buffer (25 mM HEPES, pH 7.5, 10 mM MgCl_2_, 150 mM NaCl, 0.5 mM EGTA, 20 mM β-glycerophosphate, 0.5% Nonidet-P40, 10% glycerol, and phosphatase- and protease inhibitors. The lysates were incubated with beads carrying recombinant GST-RBD protein and rotated for 1 hour at 4°C. After washing with lysis buffer, activated Ras pulled down by the GST-RBD beads was measured by Western blotting. Western blots were quantified using ImageJ and the level of Ras activation was determined by calculating the ratio of Ras identified in GST-RBD pull downs (GTP-RAS) divided by the amount of Ras detected in total cell extracts.

### RT-PCR

Parental and resistant cells were lysed and total RNA was extracted as previously published (Duffy et al., 2014). Briefly a RNeasy kit (Qiagen, UK) was used to extract total RNA and cDNA was produced using QuantiTect Reverse Transcription kit (Qiagen) following manufacturer’s instructions. An ABI 7900HT Real Time PCR System (Applied Biosystems) and TaqMan reagents (Applied Biosystems MA, USA) were used according to the manufacturer’s protocol. Gene expression was normalized to the expression of β-actin (Assay I.D: 4326315E) with P0 (RPLP0, Assay I.D: 4310879E) as a second endogenous control. Gene assays used were: MST1 (STK4, Hs00178979_m1*), MST2 (STK3, Hs00169491_m1*), LATS1 (Hs01125523_m1*), LATS2 (Hs00324396_m1*) and YAP1 (Hs00902712_g1*). Biological duplicates were generated for all samples at all time points. Technical replicates for every sample and time point were also performed

### Patient Samples and Immunohistochemistry

Patients with metastatic melanoma containing BRAF^V600E^ mutation (confirmed by genotyping) were enrolled on clinical trials for treatment with a Vemurafenib (BRAF inhibitor) or combined Vemurafenib and dabrafenib + trametinib (MEK inhibitor) and information about the study is included in Frederick et al (Frederick et al., 2013). Patient were consented for tissue acquisition per Institutional Review Board (IRB)-approved protocol (DFCI 11-181). Tumor biopsies were conducted pre-treatment (day 0), at 10-14 days on treatment and/or at time of progression. Formalin-fixed tissue was analyzed to confirm that viable tumor was present via hematoxylin and eosin (H&E) staining. Responses were determined according to RECIST, version 1.1.

Patient tumors were fixed in 10% neutral buffered formalin as previously described (Frederick et al., 2013). Briefly, tissue was embedded in paraffin, and sectioned at five microns. Deparaffinized and rehydrated sections were subjected to epitope retrieval in 10 mM Tris-EDTA buffer pH 9.0 and blocking in 3% BSA in TBST (Tris pH 7.6, 0.05% Tween-20). Sections were incubated with MST2 (STK3) antibody (Epitomics, clone EP1466Y, cat# 1943-1) for 1 hour at RT. After peroxidase block in 3% H_2_O_2_, HRP-labeled anti-rabbit secondary antibody (Dako EnVision, K4003, RTU) was applied for 30 minutes. Slides were developed with DAB+ (Dako K3468) and counterstained with hematoxylin (Vector H-3401) prior to dehydration and mounting. Stained slides were scored by 2 expert, blinded pathologists for positivity and intensity.

### Statistics

Experiments were repeated at least times unless indicated. Graphs were generated using Excel, error bars show standard deviation (SD).

## Data availability

This study includes no new data deposited in external repositories. Proteomics data used in this study have been published before (Quinn et al., 2021, Novacek et al., 2020) and are available in PRIDE repository, access numbers PXD018903 and PXD018905.

## Acknowledgements

We thank the patients that donated the tumour samples used in this study. We thank Claudia Aura González (UCD, Conway Institute) for helping to score IHC samples. This work was supported by Science Foundation Ireland under grant numbers 18/SPP/3522 and 14/IA/2395 awarded to WK, and CDA 15_CDA_3495 to DM.

## Author contributions

All authors contributed to the writing of the final manuscript. David Romano: conceptualization; investigation; writing-original draft and figures; data analysis. Lucia Garcia: investigation; data analysis; David Duffy: investigation; data analysis. Keith T. Flaherty: data analysis. Dennie T. Frederick: investigation; data analysis. Walter Kolch: supervision; conceptualization; writing original draft; project administration. David Matallanas: Conceptualization; supervision; writing original draft; writing review and editing.

## Appendix

### Supplementary figures

**Supplementary figure 1.**
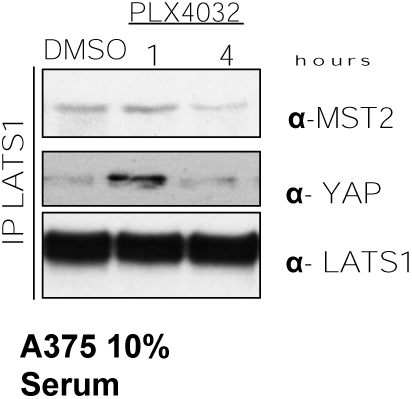
BRAF inhibition regulates the MST2 pathway. A375 cells were treated with DMSO or PLX4032 (3µM) for the indicated times. LATS1 was immunoprecipitated using specific antibody and interaction with MST2 and YAP was detected by western blot with indicated antibodies.

**Supplementary figure 2.**
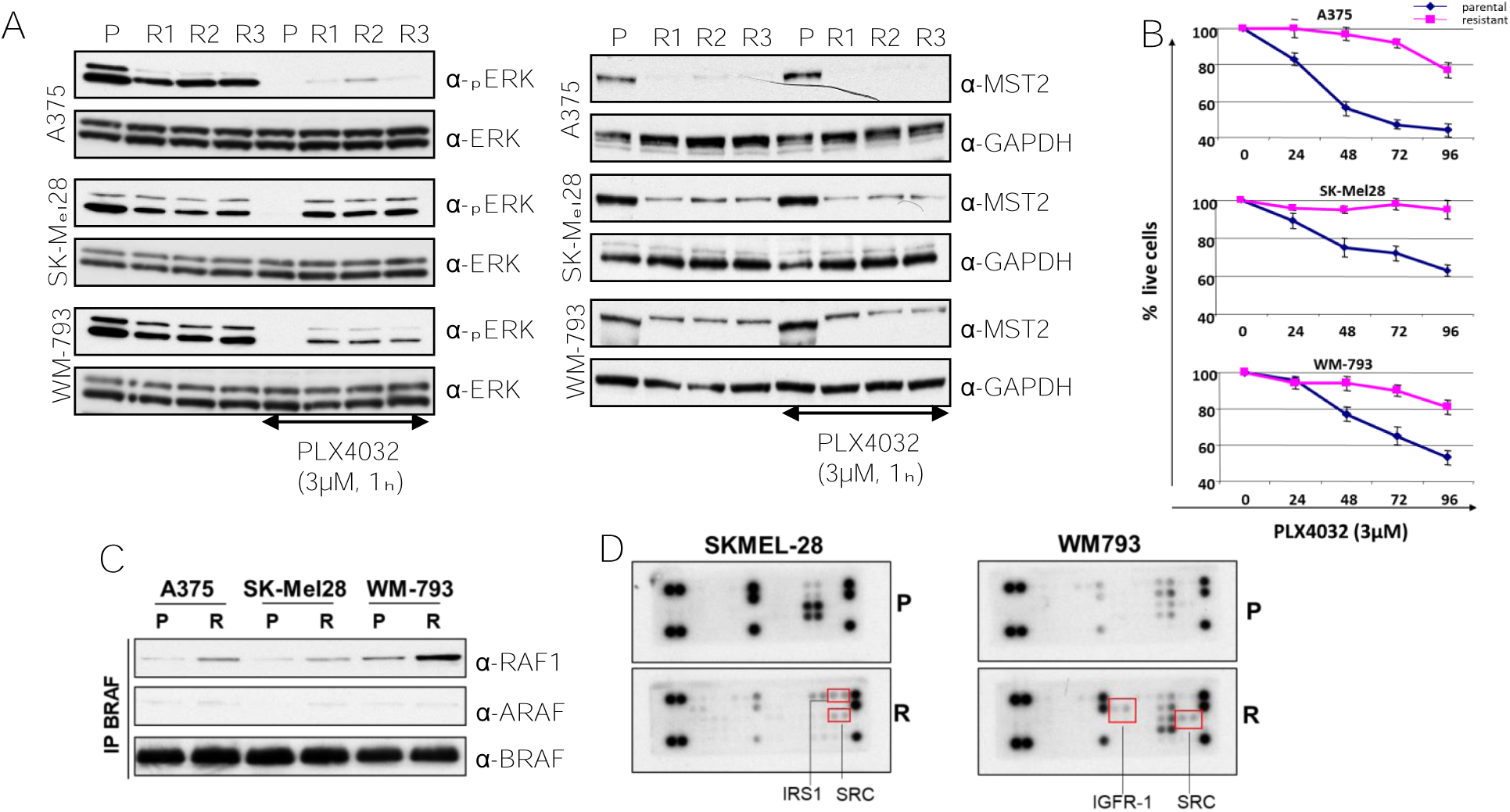
Mutant BRAF melanoma cells resistant to PLX4032 cells rewire ERK and MST2 pathways. A) Parental (P) A375, SK-Mel28 and WM-793 cells clones (R) grown without PLX4032 for 48 hours were treated with PLX4032 (3 µM) for 1 hour and ERK activation was determined by immunoblotting with the indicated antibodies. B) Cells were grown in 3 μMPLX4032, and live cells were counted at the indicated timepoints using a cell cytometer. C) Parental and resistant cells were lysed and the endogenous BRAF was immunoprecipitated. BRAF interaction with ARAF and RAF1 was determined by immunoblotting with the indicated antibodies. D) Parental (P) and resistant SKMEL-28 (left) and WM793 (right) (R) cells were lysed and cell extracts were incubated with PathScan® RTK Signaling Antibody Arrays. Red boxes show differentially regulated proteins.

**Supplementary Figure 3.**
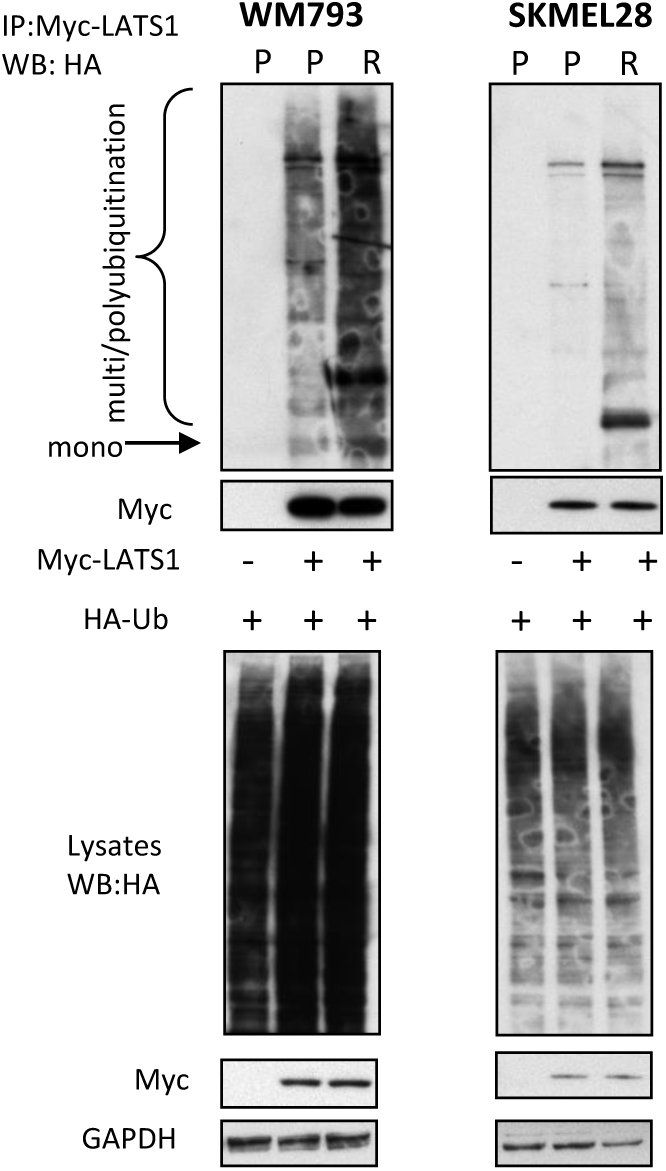
LATS1 exhibit increased ubiquitination and association with E3-ubiqitin ligases in resistant melanoma cells. Parental and resistant A375, SK-Mel28 and WM-793 cell lines were transfected with HA-ubiquitin (1 μg) alone or co-transfected with Myc-LATS1 (1 µg) and lysed 48 hours after transfection. Cell extracts were IP with Myc-tag antibody beads and ubiquitination of precipitated LATS1 was determined using HA antibody. Expression of proteins was determined in cells extract by immunoblotting with specific antibodies

**Supplementary Figure 4.**
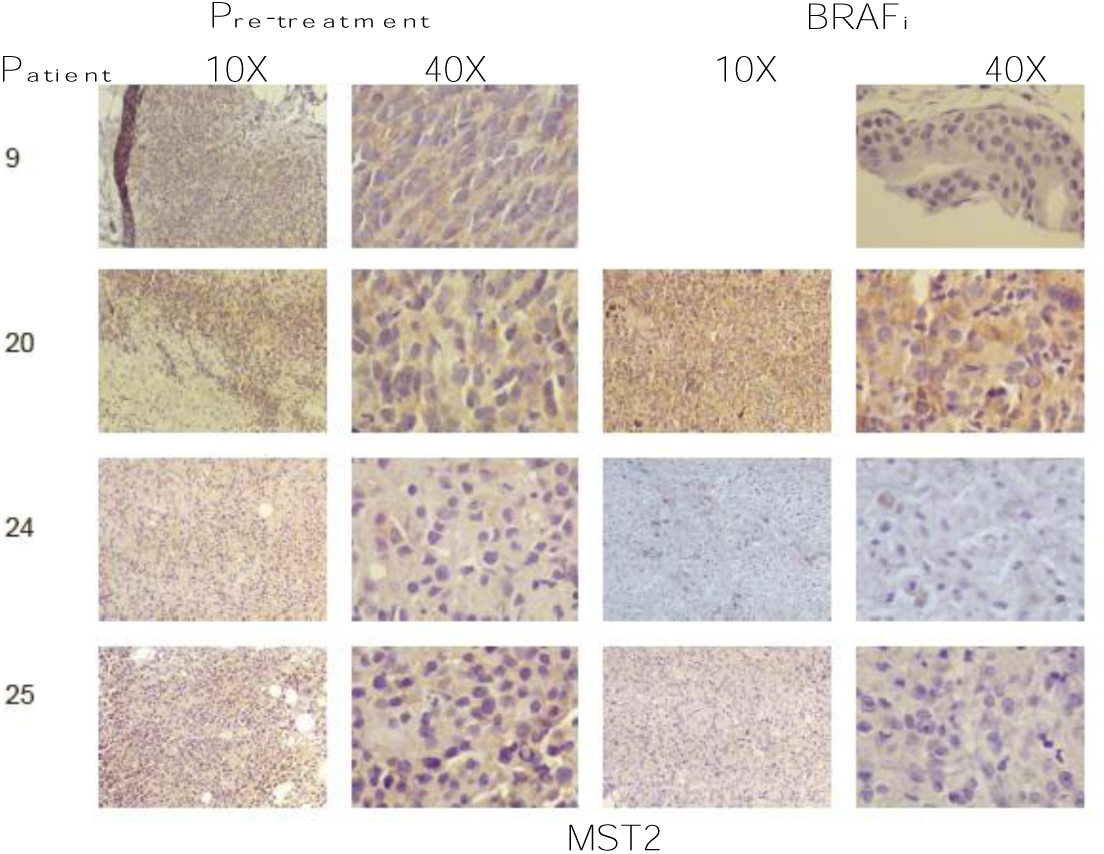
MST2 expression is decreased in patients with metastatic melanoma treated with BRAF inhibitors. Immunohistochemistry (10x and 40X magnification) for the MST2. Tumours biopsied from the indicated patients were stained with haematoxylin and eosin. The figure shows expression of MST2 in patients 20 and 24 and before treatment with BRAF inhibitor alone (pre-treatment) and after treatment (BRAFi) and 9 and 25 that were treated with BRAF and MEK inhibitors (BRAFi).

**Supplementary Figure 5.**
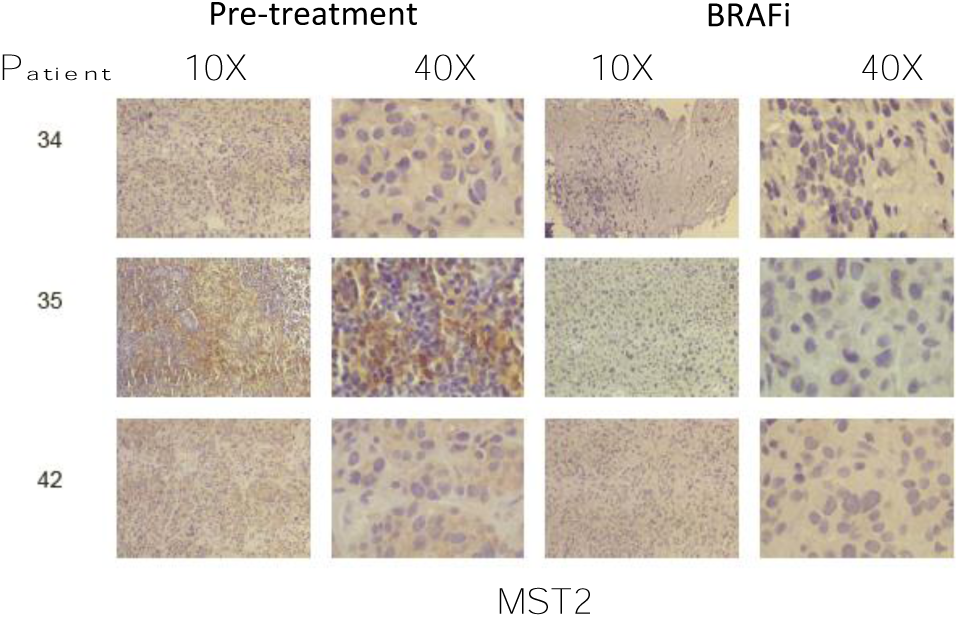
MST2 expression is decreased in patients with metastatic melanoma treated with BRAF inhibitors. Immunohistochemistry (10x and 40X magnification) for the MST2. Tumours biopsied from the indicated patients were stained with haematoxylin and eosin. The figure shows expression of MST2 in patients 34 and 35 and 42 before treatment with BRAF inhibitor (pre-treatment) and after treatment BRAFi and MEKi.

